# Opportunities to improve goat production and food security in Botswana through forage nutrition and the use of supplemental feeds

**DOI:** 10.1101/2023.12.04.569865

**Authors:** Andrew S. Cooke, Honest Machekano, Javier Ventura-Cordero, Aranzazu Louro-Lopez, Virgil Joseph, Lovemore C. Gwiriri, Taro Takahashi, Eric R. Morgan, Michael R. F. Lee, Casper Nyamukondiwa

## Abstract

Goats fulfil a central role in food security across Africa with over half of households owning or rearing goats in rural areas. However, goat performance is poor and mortality high. This study assessed the nutritional quality of commonly used feeds and proposes feed-baskets to enhance goat nutrition and health. Feeds were collected from 11 areas within the Central District of Botswana, and macronutrient analyses were conducted, including crude protein, fibre fractions, ash, and metabolizable energy (ME). Forage nutrition was compared across seasons and soil types. Additionally, seasonal supplementation trials were conducted to evaluate consumption rates of various supplements, including crop residues, pellets, Lablab purpureus, and Dichrostachys cinerea. Each supplement was provided ad libitum for a 24-hour period, and consumption rates determined. Findings revealed significant differences in nutrition among various feed sources, across seasons, and in relation to soil types (p < 0.001). Consumption rates of supplements were higher during the dry season, possibly due to reduced forage availability. Supplement consumption rates varied across supplements, with crop residues accounting for approximately 1% of dry matter intake, compared to up to 45% for pellets, 13% for L. purpureus, and 15% for D. cinerea. While wet season feed baskets exhibited higher ME values compared to dry-season feed-baskets, the relative impact of supplementation was more pronounced during the dry season. These results highlight the potential for optimizing goat diets through improved grazing and browsing management, especially during the reduced nutritional availability in the dry season.

## 1 Introduction

Across Sub-Saharan Africa (SSA), goats play vital nutritional, socio-economic, and cultural roles. This is especially true in rural communities where more than half of households own or rear goats in some capacity (Manirakiza et al., 2020). The goat population in Botswana is distributed across the country and is estimated to include approximately 1.4 million head (Mataveia et al., 2021), nearly exclusively reared by smallholders (Burgess, 2005), making it the most popular form of livestock (Bolowe et al., 2022). Goats contribute to income, food, and nutritional security through their ability to convert and store nutrients from low-value forage (graze and browse), fodder, industrial by-products, and biomass waste streams, which would otherwise be inaccessible to humans, and convert them into meat and milk. In Botswana, 29% of the population is reportedly undernourished and this appears to be increasing amidst climate and biotic shocks (World Bank, 2019a). Conversely, food insecurity (lack of available food) is slightly better than the SSA average with a rate 50.8% in Botswana compared to the SSA mean of 59.5% (World Bank, 2019b, 2019c). This disparity suggests that nutritional quality is an issue, which could be improved by greater access to meat and milk from livestock for the most vulnerable.

Goat production is predominantly extensively managed through communal rangeland forage grazing during the day and overnight kraaling, i.e. protective enclosure using thorn brush or other fencing (Walker et al., 2015). Agropastoral communal forage grazing in the central region is supported by hardveldt open bush savanna dominant on low fertility ferric luvisol sandy soils and moderately low fertile sandy loams (Pule-Meulenberg and Dakota, 2009). Typical rangeland goat production systems consist of relatively small household goat herd sizes (mean 21 goats per household), with a low off-take rate of 7.3% and a high mortality rate of 23.3% (Statistics Botswana, 2017). The most commonly cited reasons for owning goats are for cash (84%), followed by meat (58%), and milk (42%) (Bolowe et al., 2022; Monau et al., 2017). Therefore the financial benefits of rearing goats fall into two main categories, cash and insurance (Gwiriri et al., 2023; Kaumbata et al., 2020). The selling of meat, milk and live goats can be an important form of household income. Goat ownership can also provide resilience through the ability to sell or slaughter an animal in times of hardships. Nsoso et al. (2004) reported that farmers in Botswana generally opted not to sell stock regularly, but to use goats as a safety net or insurance, selling only when financial needs necessitated.

Broadly, Botswana has two distinct seasons, the wet season (summer and autumn - November to April) and dry season (winter and spring - May to October) and the quantity and quality of fodder varies with the seasons (Figure 1) (Naumann et al., 2017; Setshogo et al., 2011). Rainfall in the wet season aids plant growth, especially in herbaceous species, leading to a relative abundance and diversity of forage, with preferential nutritional profiles. During the wet season, goats are typically shepherded to grazing land in the day where they can consume a mix of browse, herbaceous plants, and pasture. At night, they are enclosed in a kraal (to prevent them from consuming crops and to prevent theft and predation) typically with little or no access to food or water. In the dry season goats roam more freely, predominantly on browse species, and are often not kraaled at night (highlighting that kraaling may predominantly be to protect crops). During the dry season, herbaceous plants significantly die back and forage availability skews towards browse species (Omphile et al., 2005), creating a shortage of feed and drop in nutrition availability and quality. The high costs of commercial supplementary feeds limit farmers’ ability to mitigate this. Nutritional assessment of alternative low cost, locally available supplementary feeds in arid environments thus aids in appropriate choices and utilization of the available feed resources for dry season strategic supplementation to alleviate nutritional deficiency related problems in goats (Aganga and Autlwetse, 2000).

**Figure 1.**
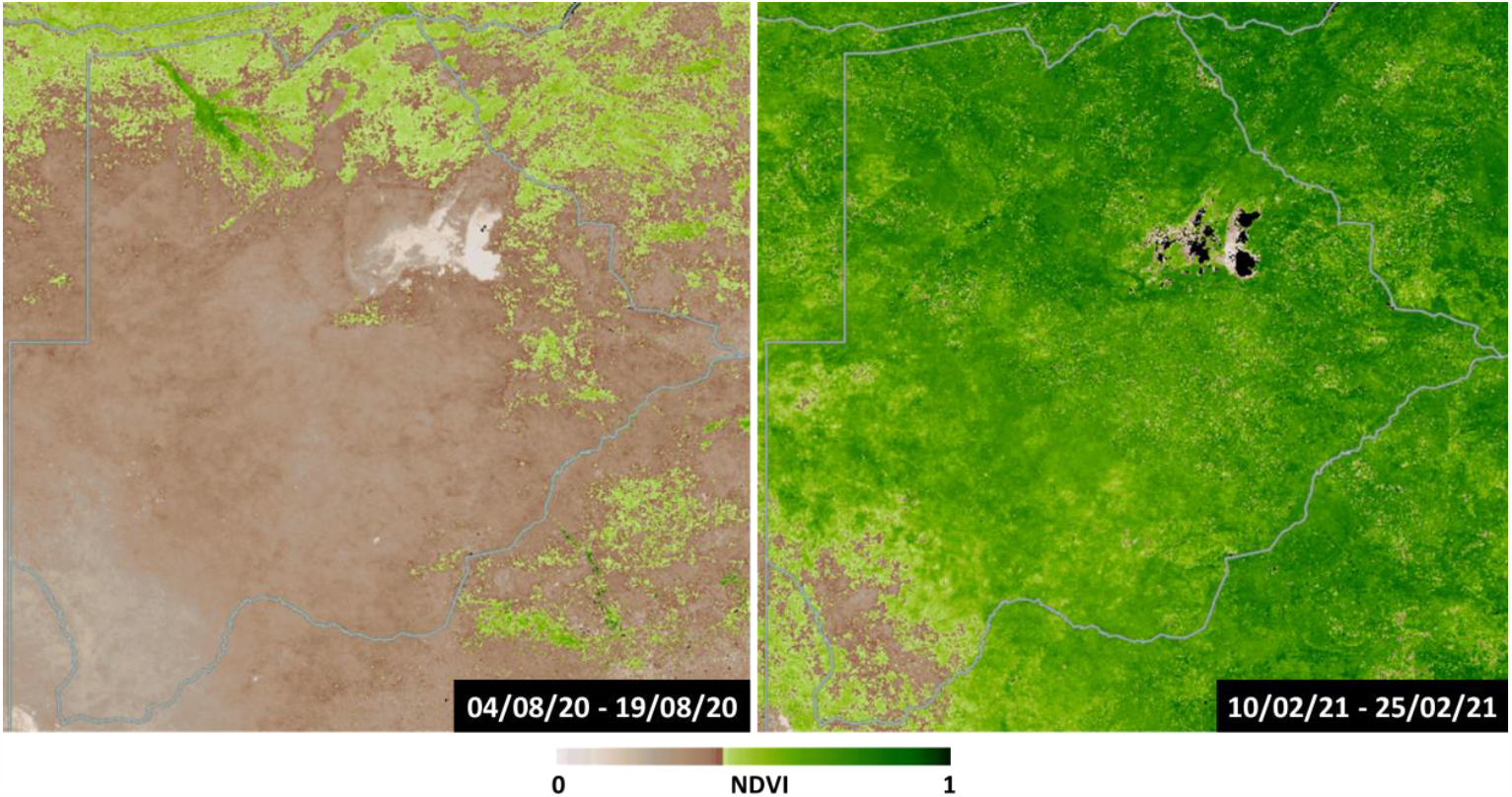
Normalised Difference Vegetation Index (NDVI ) maps of Botswana during the dry season (left) and wet Season (right). Maps are 16-day NDVI averages. Data taken from NASA Worldview (NASA, 2022).

The potential of goat enterprises has triggered several SSA governments to initiate policies that encourage investment in improving small stock-production to reduce poverty while simultaneously improving food and nutritional resilience. The government of Botswana has committed significant financial resources in small ruminants, particularly goats, through programmes such as the Livestock Management and Infrastructure Development (LIMID) program (Ministry of Agriculture, 2019) and the Remote Area Dweller Program (RADP) (Ministry of Local Government and Rural Development, 2009). Despite such investments, the productivity of goats in Botswana and SSA at large remain low due to poor nutrition, disease (e.g., gastrointestinal nematodes), and abiotic stress (e.g., frequent droughts), as well as the combined effects of such factors (Monau et al., 2017). Thus, whilst productivity is dependent on several factors, it is underpinned by optimal nutrition and disease control. By extension, improving the health and productivity of individual goats and herds could improve the resilience of these households and communities through associated household economic return or nourishment.

The objectives of this study were to:

1. Quantify the nutritional profile of cultivated and naturally available forages and feeds in the Central District of Botswana.
2. Assess the potential consumption and nutritional contribution of dietary supplements, currently used by farmers, for goat nutrition during periods where animals are kraaled.
3. Use the information obtained from objectives 1 and 2 to develop and assess theoretical feed-baskets for both the dry and wet seasons to optimise nutrition availability and quality based on available resources.

## 2 Methods

### 2.1 Forage collection and analysis

A variety of forage samples (*n =* 244) were collected across the Central District of Botswana between January 2020 and October 2021. Samples came from 21 farms/smallholdings, spanning 11 villages (Lecheng, Maape, Mhalapitsa, Mogorosi, Paje, Palapye, Pilikewe, Radisele, Ramokgnami, Serowe, Thabala) (Figure 2). Forages were selected based on farmer recollection of goats consuming them and/or physical evidence of goat grazing. The one exception to this was *Viscum* spp. which whilst not reportedly used by farmers in this study, has been reported to be used elsewhere (Madibela et al., 2000) and shows some promise as a supplement (Madibela et al., 2010; Moncho et al., 2012). Farms were classified by their underlying soil type of either ‘hardveld’ (rocky) or ‘sandveld’ (sandy) (Panagos et al., 2011). Collection dates were recorded allowing for samples to be designated as from either the dry season or wet season. Where possible species or genus information was recorded. Additionally, forages were given one of three classifications:

**browse** – plants with hard stems such as woody trees and shrubs.

**herbaceous** – non-woody species with soft stems, such as grasses and forbs.

**pasture** – This refers to flat and low-lying plains, dominated by grasses. Such areas are often under communal livestock grazing. Samples designated ‘pasture’ were not speciated and were general cuttings of a quadrat within this area and were thus typically mixes of herbaceous species.

**Figure 2.**
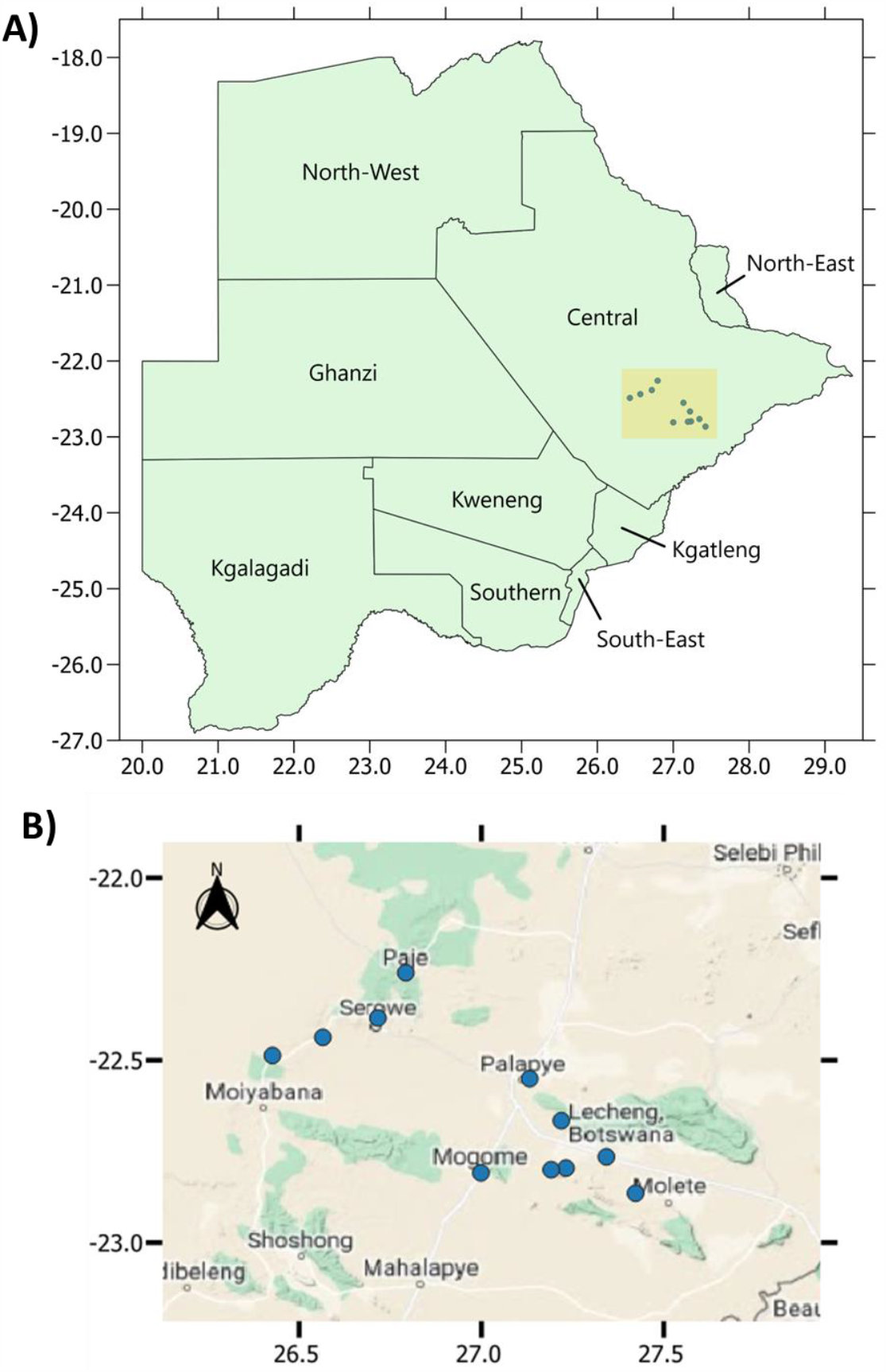
A) Map of Botswana including districts. Approximate study area highlighted in yellow with individual locales in blue dots. Axes refer to latitude and longitude. Map created using QGIS 3.26.1 (QGIS, 2022). B) Approximate location of sites. Axes refer to latitude and longitude. Map created using QGIS 3.26.1 with basemap obtained through Google Maps (Google, 2021; QGIS, 2022).

For herbaceous plants, the aerial parts (stems, leaves, stolons, flowers, fruits and/or seeds) were collected by cutting the plant stem from its base. For browse, only the browsed aerial plant parts were collected; depending on the plant species and associated browsing preference of the goats, other specific plant parts such as tender shoots, pods or flowering parts were specifically collected particularly for *Dichrostachys cinerea* and different *Acacia* species. Over repeat visits, samples were collected from the same grazing area unless farmers indicated otherwise, then the new site would be sampled. During the dry season, plant supplements used by farmers were collected directly from the feeding troughs or from the storage areas. In each case, sub samples from different sampling points were mixed to make a compound sample for each type of feed.

### 2.2 Chemical analyses

Samples were weighed before being oven-dried (60°C for 48h) weighed again, vacuum packed and shipped to the UK where they were freeze dried to meet import and quarantine requirements and then ground to < 2 mm particle size for nutritional analysis. Loss on ignition was conducted (0.5 g, 540°C, 6 h) to determine ash content. Crude protein (CP) was determined as 6.25 times nitrogen content, as determined by the Dumas technique (Ebeling, 1968). Three fibre fractions were determined, neutral detergent fibre (NDF), acid detergent fibre (ADF), and acid detergent lignin (ADL) (Goering and Soest, 1970). Metabolizable energy (ME) concentrations (ME MJ kg^-1^ DM) were estimated as per Minson (1984): *ME =*10.738 + 0.161CP(%) – 0.131ADF(%). This predictive equation was chosen as it was derived from results of five tropical (*Digitaria* spp.) grasses and had recently been determined by Lwin et al. (2022) to have the best predictive value (of 23 tested) for ME of *Sorghum bicolor*, showing that the equation’s accuracy stood up across species.

Variations in forage ME concentration were compared across three plants found to be abundant across time and space: the trees *Boscia* spp. and *Terminalia* spp., and the hemi-parasitic mistletoe shrub *Viscum* spp. Two ANOVAs were conducted, the first comparing ME concentrations of the three species across time (wet season and dry season) and the second across soil type (Hardveld and Sandveld). Post-hoc Tukey testing determined differences between groups. Significance was set at α = 0.05. Analyses were performed in R and R Studio (R Core Team, 2021; R Studio Team, 2020).

### 2.3 Supplementation trials

Supplementation trials were conducted at two timepoints, the first during the wet season at the end of March (30/03/21 to 31/03/21) and the second in the dry season at the end of July (27/07/21 to 30/07/21). During the wet season, trials were conducted across eight farms: four used a crop residue (mainly maize stover (*Zea mays*) with some salt and miscellaneous plant material) and four used commercial goat pellets (Lubern Voere^®^, Hartswater, South Africa). The pellets’ composition on the label was stated as 12.9% protein, 0.7% urea, 1.5% fat, 12.9% fibre, 0.3% phosphorus and moisture content of 12.9%. During the dry season, four different supplements were tested: crop residues (as previously), *Lablab purpureus* beans, crushed pods of the leguminous tree *Dicrostachys cinerea* and commercial pellets, each replicated four times (four farmers). These supplements were chosen based on our presurvey results in the areas and anecdotal evidence observed during other research activities as representing the most commonly available and accessible type of supplements used by farmers in these areas. Supplement samples underwent nutritional analysis as per forage samples. Moisture content was calculated pre- and post- trial so that moisture loss could be accounted for in consumption rates and moisture/dry matter analysis then performed in the laboratory (60°C for 48 hrs) to enable DMI determination.

Each trial was conducted in a similar manner: A weighed ration of the supplement (Table 1) was provided to the flock in the afternoon (when the goats were coming back to the kraal for the night) for the goats to consume until noon the next day (approx. 19hrs). The supplement was therefore available after access to the basal diet, prior to kraaling, which constituted predominately herbaceous plants and browse during the dry season and pasture and browse during the wet season. No other feeds were available to the goats once kraaled. After this period, any remaining supplement was re-weighed (when the goats were released the next day) to assess how much had been consumed at herd level, which was then adjusted for moisture loss and consumption on a per animal basis calculated. However, as individual animal weights were not known, and each flock had a different composition, an adjustment factor was imposed. Goats were categorised into one of four categories: (1) Adult female (2) Adult male (3) Female kid (4) Male kid, with kids being < 1 year old. Mean weights for each of these categories were taken from (Katongole et al., 1996) and the mean of those four weights (25.35 kg) considered to be the weight of a typical goat (hereon referred to as a ‘goat unit’). The mean weights of each category (as per Katongole et al. (1996)) was then calculated relative to that value, providing an adjustment factor (Table 2). These adjustment factors were then applied to the known group composition to allow for consumption to be calculated based relative to ‘goat units’. Target DMI for goats was considered as 4% of liveweight (Freking and McDaniel, 2016), equating to 1.01 kg per goat unit per day.

**Table 1.**
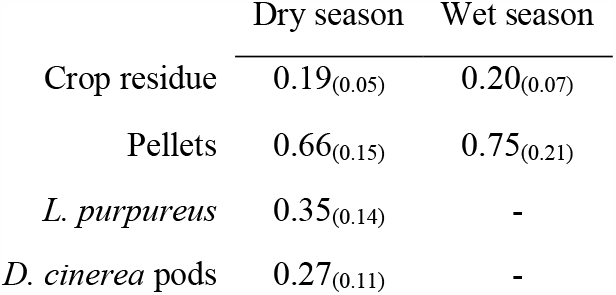
Provision of supplements (kg, mean, on a per goat unit basis) of each supplement for the dry and wet season trials. Subscripted number in brackets is standard deviation.

**Table 2.** Adjustment factors to standardise consumption across different goat types. Typical weights taken from Katongole et al. (1996).

| Category | Mean weight (kg) | Goat units |
| --- | --- | --- |
| Female adult | 28.99 | 1.14 |
| Male adult | 33.39 | 1.32 |
| Female kid | 19.64 | 0.78 |
| Male kid | 19.23 | 0.76 |

### 2.4 Feed-basket formulation

Forage nutrition data and supplement trial results were used to assess numerous theoretical feed-baskets available to the goats. For the basal diet (browse species, herbaceous species, pasture, that had *n* > 1 samples) and supplements, mean ME and CP concentrations were taken for each season (where available). Each feed-basket comprised a basal diet (Herbaceous and Browse, during the dry season; and Pasture and Browse, during the wet season) and supplementation (including a control with no supplementation). Basal diets were a varying ratio of the naturally available forage types at that time. During the wet season goats graze predominantly on the abundant pasture forages and on browse, consequently the basal diet was a ratio of the two from 100:0 to 0:100 in steps of ±20. For the dry season, the pasture plants die off, though some herbaceous species persist and can make up around 10- 25% of total diet, thus the basal diet for this period was comprised of herbaceous and browse plants at ratios from 25:75 to 0:100 in steps of ±5. The contribution of the basal diet to the overall diet was adjusted to make way for supplementation. Supplement inclusion rates were set at the level of intake (as a proportion of DMI targets) observed in the supplement trials. *Viscum* spp. was also added as a theoretical supplement at a rate of 20.0% as per Madibela and Jansen (2003), despite not being tested directly in the feed trials. For each theoretical feed-basket (wet season: *n* = 24, dry season: *n* = 36) the ME and CP concentrations of the formulated feed baskets were then calculated, as well as the ratio of CP to ME (CP:ME).

## 3 Results

### 3.1 Forage nutrition

There was a statistically significant difference in ME concentrations across the three forages *Boscia* spp., *Terminalia* spp. and *Viscum* spp. (*F* = 214.1, *p* < 0.001) (Table 3). Seasonal differences in nutritional composition were observed across the entire sample pool (*F* = 31.0, *p* < 0.001), with samples collected in the wet season yielding the highest ME concentrations (Figure 3). However, this was less apparent intra-species with Tukey testing only showing a significance between season for *Terminalia* spp., though dry season ME concentrations were lower than in the wet season for both *Boscia* spp. and *Viscum* spp. Across these three species there was also a significant difference in ME based on the underlying soil type (*F* = 27.4, *p* < 0.001), with Sandveld soils yielding higher median ME concentrations than Highveld (Figure 4) for all species. However, within each species, Tukey testing did not reveal a significant difference between soil types.

**Table 3.**
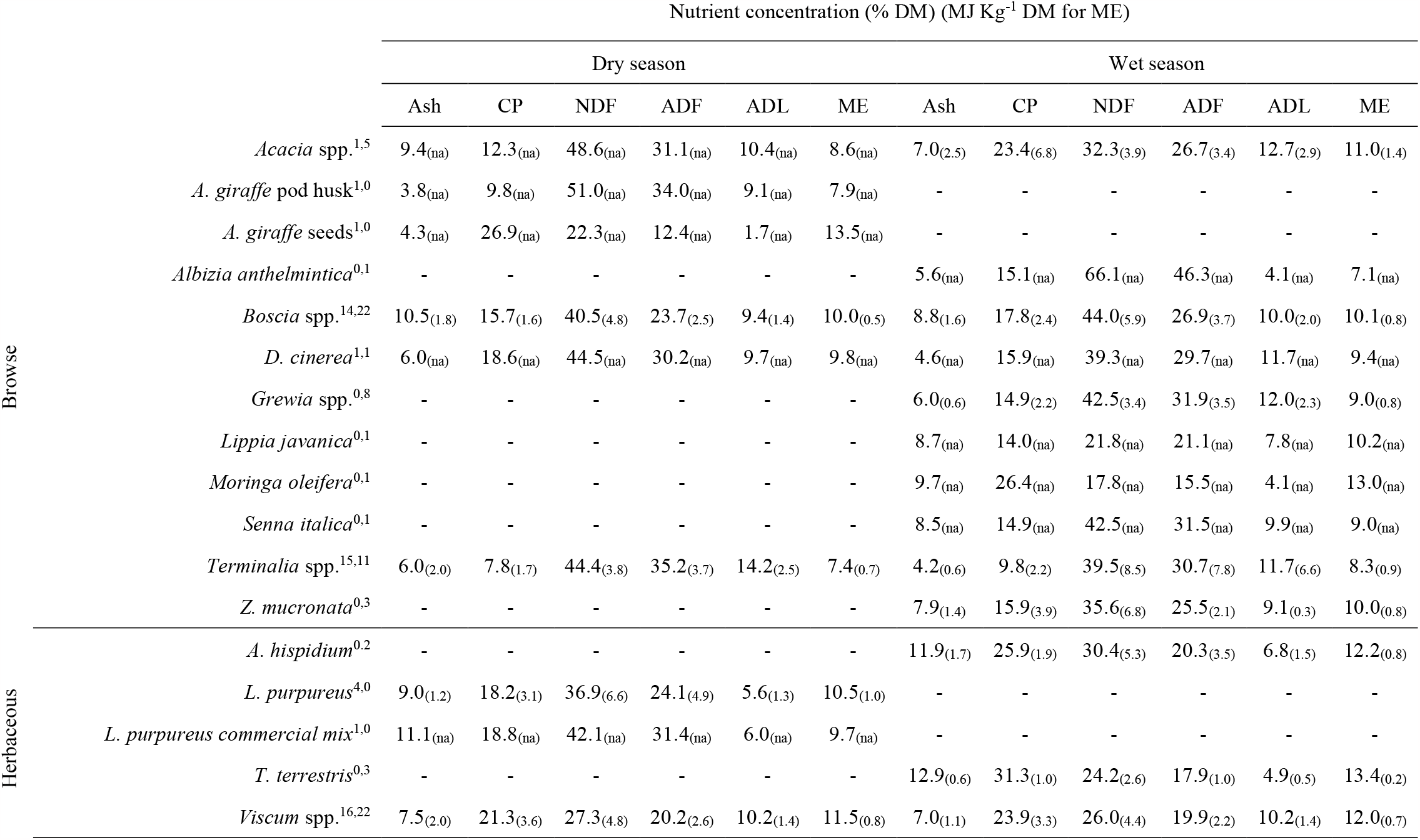
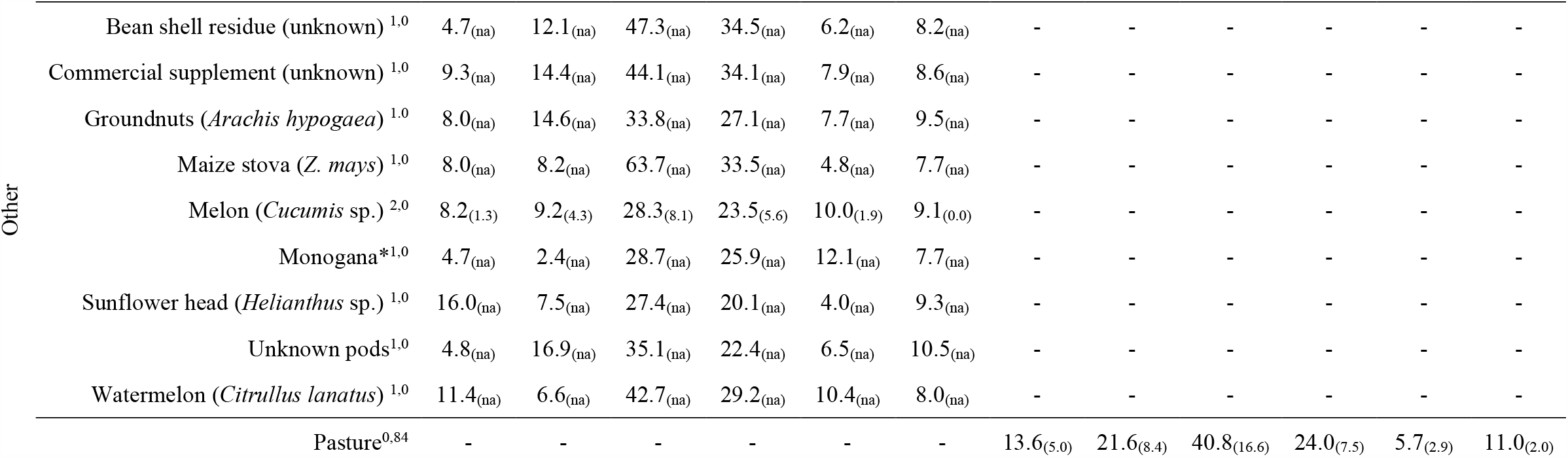
Nutritional profiles of browse plants, herbaceous plants, and pasture samples during the dry season and wet season. Concentrations are expressed as % DM, except for ME which is expressed as MJ kg-1 DM. Superscript numbers after species names signify sample size (*n*) for the two seasons respectively. Subscript after values represent standard deviation (where available). See text, methods section, for abbreviations.

**Figure 3.**
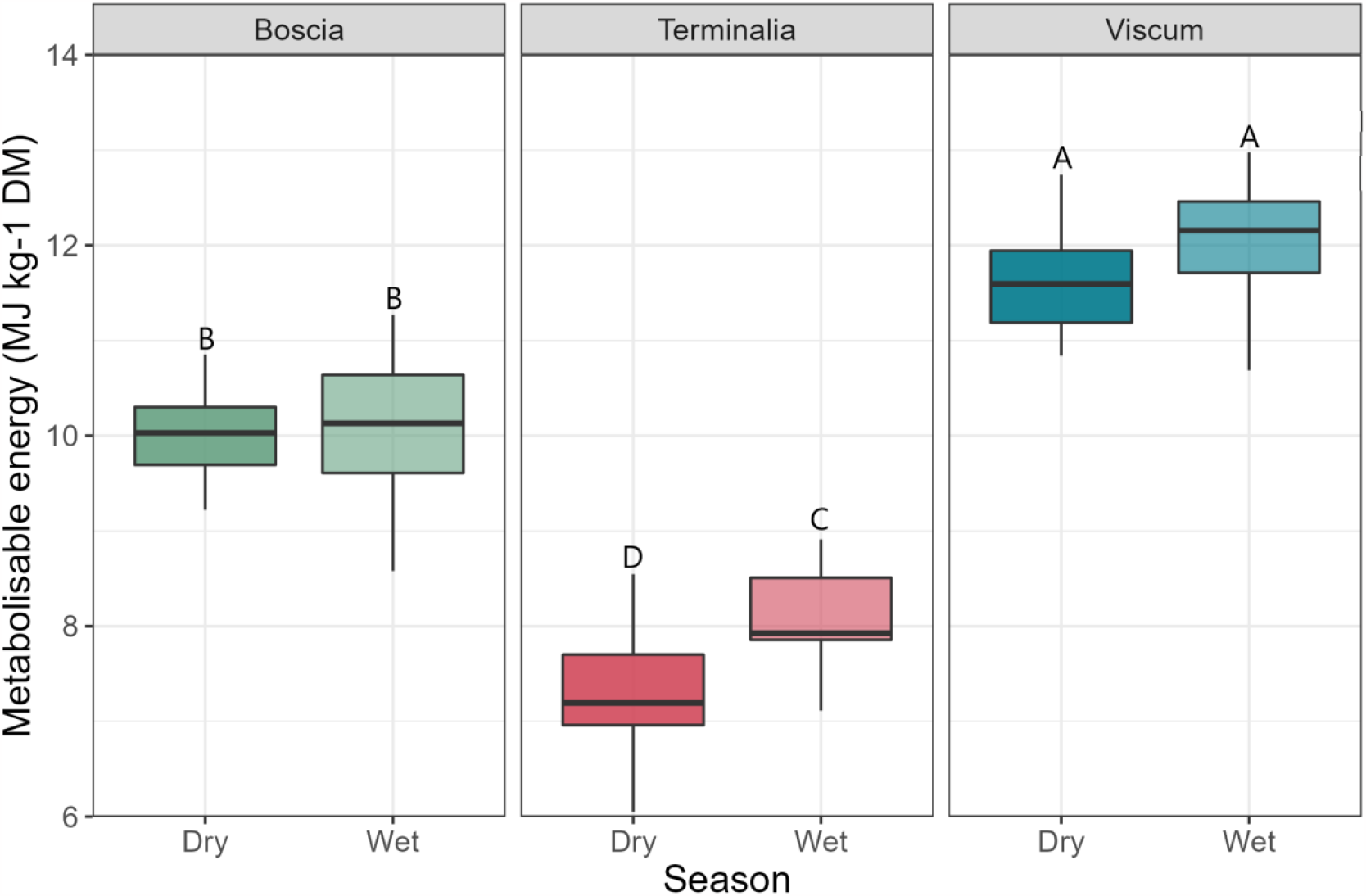
Metabolizable energy concentrations (MJ kg^-1^ DM) of *Boscia* spp., *Terminalia* spp., and *Viscum* spp. between the dry and wet seasons. Boxplots sharing letters are not significantly different to one another.

**Figure 4.**
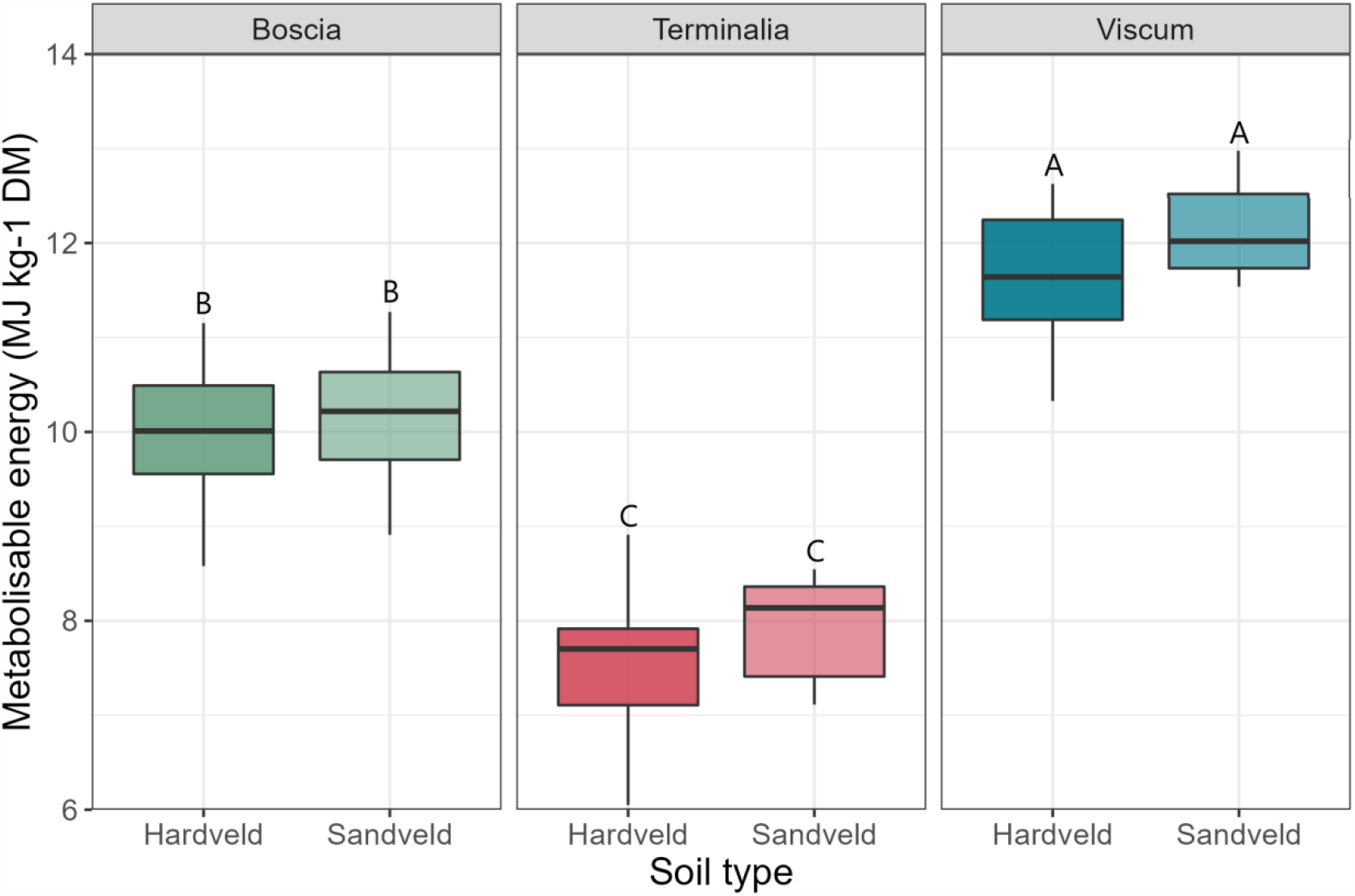
Metabolizable energy concentrations (MJ kg^-1^ DM) of *Boscia* spp., *Terminalia* spp., and *Viscum* spp. between samples obtained from Hardveld soils and Sandveld soils. boxplots sharing letters are not significantly different to one another.

### 3.2 Supplement trials

The nutritional profile of supplements varied greatly (Table 4). Crop residues had the lowest CP, ADF and ADL concentrations. Pellets and *L. purpureus* had middling profiles in all regards, CP was above minimum requirements (5-7%) (Lazzarini et al., 2009; Pugh, 2020), but lower than optimal (15-17%) (Salah, 2015). NDF:ADF ratios were around 4:3. *Dichrostachys cinerea* stood out in terms of high CP concentrations, low ash content, and low ADF.

**Table 4.** Nutritional profile of supplementary feeds used in feeding trials. Concentrations are expressed as % DM, with the exception of ME which is expressed as MJ kg-1 DM.

|  | Ash | CP | NDF | ADF | ADL | ME |
| --- | --- | --- | --- | --- | --- | --- |
| Mixed crop residue | 10.9 | 5.3 | 44.0 | 25.8 | 4.5 | 8.2 |
| Pellets | 10.3 | 12.4 | 42.3 | 26.4 | 6.3 | 9.3 |
| *L. purpureus* | 7.8 | 10.8 | 47.6 | 31.5 | 7.3 | 8.4 |
| *D. cinerea* pods | 4.3 | 16.5 | 40.4 | 27.3 | 8.9 | 9.8 |

Crop residue consumption rates were low across both seasons, at around 0.01 kg (10 grams) per goat unit and ≤10% of total provision (Figure 5). In the wet season trials, this equated to around 0.6% of daily DMI targets, doubling to 1.2% in the dry season (Table 5). Conversely, consumption rates of pellets were high, with herds consuming half or more of their allocation, equating to an average of 34.9% of their daily DMI target in the wet season and 44.5% in the dry season (+27.5%). Consumption rates of *L. purpureus* and *D. cinerea* pods were moderate, with goats consuming approximately half of the ration. In no cases did the total provision or availability of supplement appear to be a limiting factor to consumption.

**Table 5.** Mean percentage of target dry matter intake (4% liveweight of one goat unit = 1.01 kg) met by supplementation.

|  | Dry season |  | Wet season |  |
| --- | --- | --- | --- | --- |
|  | Mean % target DMI | S.D. | Mean % target DMI | S.D. |
| Crop residue | 1.2 | 0.5 | 0.6 | 0.4 |
| Pellets | 44.5 | 16.0 | 34.9 | 8.1 |
| *L. purpureus* | 12.5 | 3.2 | - | - |
| *D. cinerea* | 14.6 | 8.6 | - | - |

**Figure 5.**
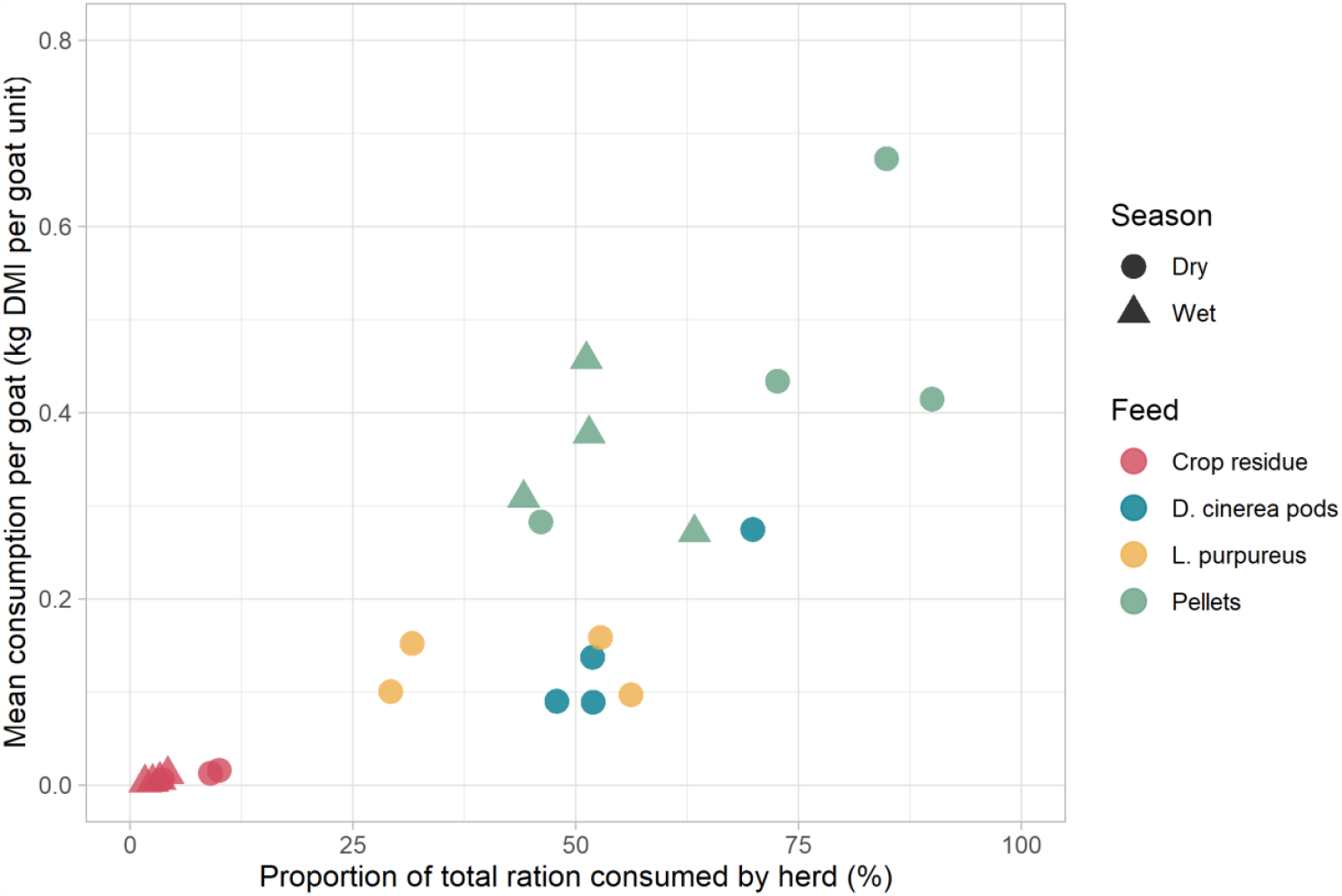
- Consumption rates of supplements during supplementation trials. Each point refers to an individual trial on one farm. Note: one result for crop residue consumption in the dry season was voided as goats spilled the feed bucket and thus quantification of consumption was not possible.

### 3.3 Feed-baskets

Wet season feed-baskets typically had higher ME and CP concentrations than dry season feed-baskets (Table 6 and Table 7). Both the highest and lowest CP:ME ratios were observed in the wet season feed-baskets (Table 8) and these were predominantly driven by the basal diet (pasture: browse ratio), as opposed to supplementation. Supplementation with crop residue had little impact on ME and CP concentrations, due to its low inclusion level. Pellets had no strong effect on ME in the dry season but had a small effect in the wet season. Notably, pellets had a large negative impact on CP across both seasons, due to their low CP concentration and high intake rate. *L. purpureus* (dry season only) had a small negative effect on CP and to a lesser extent ME. *D. cinerea* pods had a small positive effect on ME and a small negative effect on CP. *Viscum* spp. provided moderate gains to ME across both seasons, yielding the most energy dense feed baskets. During the dry season, it marginally lowered CP, due to the high CP content of the basal diet, though for the wet season it provided a moderate increase in CP.

**Table 6.** Metabolizable energy (ME) concentrations (MJ kg^-1^ DM) of theoretical feed-baskets. Supplementation rates are derived from trial results (Table 5). Shading is relative to cell value. The table provides sufficient information to enable the reader to estimate ME concentrations of other rations of these feeds.

|  |  | Herbaceous : Browse |  |  |  |  |  |
| --- | --- | --- | --- | --- | --- | --- | --- |
| **Dry season** |  | 25:75 | 20:80 | 15:85 | 10:90 | 5:95 | 0:100 |
| None (0.0%) |  | 9.15 | 9.06 | 8.97 | 8.88 | 8.78 | 8.69 |
| Crop residue (1.2%) |  | 9.14 | 9.05 | 8.96 | 8.87 | 8.78 | 8.69 |
| Pellets (44.5%) |  | 9.20 | 9.15 | 9.10 | 9.05 | 9.00 | 8.95 |
| *L. purpureus* (12.5%) |  | 9.05 | 8.97 | 8.89 | 8.81 | 8.73 | 8.65 |
| *D. cinerea* (14.6%) |  | 9.24 | 9.17 | 9.09 | 9.01 | 8.94 | 8.86 |
| *Viscum* spp. (20.0%) |  | 9.62 | 9.55 | 9.48 | 9.40 | 9.33 | 9.26 |
|  |  | Pasture : Browse |  |  |  |  |  |
| **Wet season** |  | 100:0 | 80:20 | 60:40 | 40:60 | 20:80 | 0:100 |
| None (0.0%) |  | 10.58 | 10.38 | 10.18 | 9.98 | 9.78 | 9.58 |
| Crop residue (0.6%) |  | 10.56 | 10.36 | 10.17 | 9.97 | 9.77 | 9.57 |
| Pellets (34.9%) |  | 10.12 | 9.99 | 9.86 | 9.73 | 9.60 | 9.47 |
| *Viscum* spp. (20.0%) |  | 10.86 | 10.70 | 10.54 | 10.38 | 10.22 | 10.06 |

**Table 7.** Crude protein (CP) concentrations (% DM) of theoretical feed-baskets. Supplementation rates are derived from trial results (Table 5). Shading is relative to cell value. The table provides sufficient information to enable the reader to estimate CP concentrations of other rations of these feeds.

|  |  | Herbaceous : Browse |  |  |  |  |  |
| --- | --- | --- | --- | --- | --- | --- | --- |
| **Dry season** |  | 25:75 | 20:80 | 15:85 | 10:90 | 5:95 | 0:100 |
| None (0.0%) |  | 17.92 | 17.92 | 17.93 | 17.94 | 17.94 | 17.95 |
| Crop residue (1.2%) |  | 17.77 | 17.77 | 17.78 | 17.79 | 17.79 | 17.80 |
| Pellets (44.5%) |  | 15.46 | 15.47 | 15.47 | 15.47 | 15.48 | 15.48 |
| *L. purpureus* (12.5%) |  | 17.03 | 17.03 | 17.04 | 17.05 | 17.05 | 17.06 |
| *D. cinerea* (14.6%) |  | 17.71 | 17.72 | 17.72 | 17.73 | 17.73 | 17.74 |
| *Viscum* spp. (20.0%) |  | 17.79 | 17.79 | 17.80 | 17.81 | 17.81 | 17.82 |
|  |  | Pasture : Browse |  |  |  |  |  |
| **Wet season** |  | 100:0 | 80:20 | 60:40 | 40:60 | 20:80 | 0:100 |
| None (0.0%) |  | 17.64 | 18.33 | 19.03 | 19.72 | 20.42 | 21.11 |
| Crop residue (0.6%) |  | 17.56 | 18.25 | 18.94 | 19.63 | 20.32 | 21.02 |
| Pellets (34.9%) |  | 15.81 | 16.26 | 16.71 | 17.17 | 17.62 | 18.07 |
| *Viscum* spp. (20.0%) |  | 18.50 | 19.06 | 19.62 | 20.17 | 20.73 | 21.28 |

**Table 8.** Crude protein to metabolisable energy ratio of theoretical feed baskets (grams CP per MJ ME, dry matter basis). Supplementation rates are derived from trial results (Table 5). Shading is relative to cell value.

|  |  | Herbaceous : Browse |  |  |  |  |  |
| --- | --- | --- | --- | --- | --- | --- | --- |
| **Dry season** |  | 25:75 | 20:80 | 15:85 | 10:90 | 5:95 | 0:100 |
| None (0.0%) |  | 19.6 | 19.8 | 20.0 | 20.2 | 20.4 | 20.7 |
| Crop residue (1.2%) |  | 19.4 | 19.6 | 19.8 | 20.1 | 20.3 | 20.5 |
| Pellets (44.5%) |  | 16.8 | 16.9 | 17.0 | 17.1 | 17.2 | 17.3 |
| *L. purpureus* (12.5%) |  | 18.8 | 19.0 | 19.2 | 19.4 | 19.5 | 19.7 |
| *D. cinerea* (14.6%) |  | 19.2 | 19.3 | 19.5 | 19.7 | 19.8 | 20.0 |
| *Viscum spp.* (20.0%) |  | 18.5 | 18.6 | 18.8 | 18.9 | 19.1 | 19.2 |
|  |  | Pasture : Browse |  |  |  |  |  |
| **Wet season** |  | 100:0 | 80:20 | 60:40 | 40:60 | 20:80 | 0:100 |
| None (0.0%) |  | 16.7 | 17.7 | 18.7 | 19.8 | 20.9 | 22.0 |
| Crop residue (0.6%) |  | 16.6 | 17.6 | 18.6 | 19.7 | 20.8 | 22.0 |
| Pellets (34.9%) |  | 15.6 | 16.3 | 16.9 | 17.6 | 18.4 | 19.1 |
| *Viscum spp.* (20.0%) |  | 17.0 | 17.8 | 18.6 | 19.4 | 20.3 | 21.2 |

## 4 Discussion

The protein and energy requirements of goats will depend on a whole array of factors, both biotic and abiotic, including breed, level of performance, health status, and thermoregulation; but assuming a level of lactation (0.5 - 1 litre) or moderate body weight gain of ca. 20 g/day goats will require approximately 9.4 MJ/day and 54 g metabolizable protein (modified from AFRC, 1993; assuming q_m_ = 0.59). Assuming also a ratio of metabolizable protein : crude protein of 0.64 – 0.80 (Cannes et al., 2008) would equate to roughly 84.4 – 67.5 g CP/day plus 55 g CP/litre of milk. Of course, such values are predicted from equations using European breeds and conditions but provide a range of target intakes to assess African diets, until more detailed understanding of the protein and energy requirements of African goats under local conditions and diets is available. As such the availability of the key nutrient’s protein and energy, notwithstanding water, and micro-nutrients (which this paper does not consider), evaluated in this study from the basal diets (herbaceous plants and browse consumed prior to kraaling) were critically constraining for ME in the dry season emphasising the critical role of supplementation. Available nutrition was more favourable in the wet season, consistent with other reports from SSA (Omphile et al., 2005; Setshogo et al., 2011).

The vegetation of arid range land is dominated by browse, in the form of shrubs, bushes and sub-shrubs (van Duivenbooden, 1989), and they form an integral part of the farming system in the humid zone, particularly of west Africa (Atta-Krah et al., 1986). In terms of utilisation, browse currently play an important, albeit non-strategic role in goat nutrition, as animals under confinement in the humid zone often receive one type or the other of browse, from fallow lands or around the homestead, forming up to 25% of their diet. In the arid and semi-arid zones, browse constitute the main feed resource during the extended dry periods of the year (Le Houerou, 1980) and play a similar role in the sub-humid savannah zone. The nutritional value of browse has also been exploited in feeding systems using them as supplements to low quality tropical forages and crop residues. In general, many of the common browse species contain high levels of protein and energy in the range of 14 to 26% CP and 11 to 14 MJ of ME/kg of dry matter. In addition, they have good levels organic matter digestibility (50-60%), and contain reasonable levels of both macro and trace minerals required for efficient rumen function (Smith, 1992).

For a typical browse species identified in the current study, *Terminalia*, which made up a key component of many of the basal diets concentrations of CP and ME were low, especially in the dry season. For example, CP was only just above maintenance requirements providing ca. 78 g CP/kg DM, which is also when *Terminalia* is likely to make up a greater proportion of the diet due to lack of available grazing. Therefore, goats consuming a high proportion of *Terminalia* may be limiting their protein and energy intake. Conversely, CP and ME levels in *Viscum*, a potential supplement, were high all year round. Typically, goats do not consume *Viscum* in Botswana, predominantly due to it being a parasitic plant high up in its host trees which is difficult to reach, thus requiring harvesting by farmers. However, as *Viscum* lives on trees, including *Terminalia* and *Acacia*, this may provide an opportunity for, farmers to compensate for the lower protein and energy contents of these trees by supplementing with V*iscum* from the very same trees. Furthermore, parasitism of fruit trees by *Viscum* is a limiting factor to fruit yields and there is therefore a potentially synergy if *Viscum* could be harvested from orchards. Madibela et al. (2000) reported favourable dry matter and protein degradability of *Viscum* in Botswana. *Viscum* is also reported to have nutraceutical/anthelmintic properties (Madibela et al., 2010; Madibela and Jansen, 2003; Moncho et al., 2012), which may mitigate negative health impacts from infections such as gastrointestinal nematodes, which themselves act to reduce protein assimilation. For the supplements provided during both seasons (crop residue and pellets), intake was considerably higher in the dry season, hence goats were likely to consume supplements to mitigate nutrient/DMI deficiencies. This is consistent with feeding practices in SSA, where livestock generally depend on natural forage during the wet season and are only supplemented during the dry season. Pellets showed the potential to provide between a third (wet season) to a half (dry season) of target DMI, the main drawback being their cost and availability. Alternatively, *D. cinerea* pods, and to a lesser extent *L. purpureus*, may be a compromise, as they had favourable nutritional profiles and could make up 10-15% of DMI requirements. They are readily available and may be accessible at low cost in communal areas. Crop residues, predominately stovers, were not particularly desirable to goats as a supplement, though their precise composition was unknown and different residue mixes may vary. Despite the low CP of crop residue, the ADF concentration was favourable and high enough to meet requirements for rumen health, if little other feed was available. Crop residues may be a more available resource than other supplements and thus a more practical option for farmers practising mixed farming, who may use *D. cinerea* pods and *L. purpureus* alongside crop residues, assuming complementary and/or synergistic roles of these supplementary feeds. Low quality crop residues, such as stover, therefore should be considered as a resource to ensure rumen function, i.e. functional fibre to supplement higher quality feeds (Giger-Reverdin, 2017), or as a last resort basal feed during extreme dry periods where little other feed resources are available. Furthermore, in the event of crop-failure, which is becoming increasingly likely under the pressures of climate change, the consumption of failed crops by ruminants may be one way to ensure that resource is most efficiently utilised for food production. A constraint of the current study was that supplementation at any one farm was from a single supplement source, which may limit the potential of mixing different supplements to balance protein and energy requirements in a true feed-basket or total mixed ration approach. Of course, those rations would also consider other nutrients not evaluated here such as micro-nutrients.

Nutritional differences were apparent, albeit relatively minor, between farms on Hardveld and Sandveld soils. Results suggest that farms in Hardveld soil areas may benefit most from supplementation or other interventions. This study was conducted in a limited geographic range and thus when considering wider spatial variation across Botswana and SSA, further differences in plant nutritional composition (e.g., micro-nutrient composition, as already reflected) are expected to result from soil type and climatic differences, in line with wide ranges reported in the literature. However, the biggest factor facing productivity for crop-livestock farmers, specifically, will be dry matter yield of pasture in relation to soil fertility and rain fall. Mutali and Dzowela (1985) and Onifade and Agishi )1990) predicted native grassland dry matter yield to be between 1.1 – 3.2 t DM per ha per year. Therefore, with resources limited especially within crop-livestock systems the lower dry matter demands of goats would be significantly advantageous over cattle systems.

The ME and CP concentrations of feed baskets were lower in the dry season than the wet season, which meant that supplementation had a greater relative impact in the dry season compared to the wet season, highlighting temporal opportunities in nutritional intervention. Although not analysed in this work, it is likely that dry season forages had lower digestibility (Aganga et al., 2005) which would make it less likely for goats to meet their daily DMI requirement, thus increasing the relative value of supplementation further. Importantly, whilst the addition of a supplement of lower quality than the basal diet will lower the nutritional composition of the overall diet, that may be acceptable if it increases overall energy intake by making up for a shortfall in DMI, or availability of feed during periods of kraaling. During the dry season there is a stronger case for supplementation due to the lower availability and nutritional quality of forages and lower animal performance (Kraai et al., 2022). This could be most effectively targeted towards vulnerable individuals such as weanlings, pregnant does, or animals with suspected illness. The CP:ME ratio is an important determinant of a diets ability to support animal growth / performance and efficiency of nitrogen use. Low ratios would impair growth and performance limited by protein availability, whereas high values would reduce the efficiency of protein capture in the rumen leading to poor nitrogen use efficiency. All the reported diets had high CP:ME ratios which further highlights the limiting nature of available energy in these diets. Zhang et al. (2020) reported a reduction in nitrogen excretion and an increased nutrient utilization through improving rumen fermentation, enhancing nutrient digestion and absorption, and altering rumen microbiota in growing goats when reducing CP:ME from 11.3 to 8.69, whereas in the current study ratios ranged from 15.6 – 22.0. Although, dry season CP:ME ratios were less variable (16.8-20.7) than in the wet season (15.6-22.0), with lower wet season ratios associated with a higher ratio of pasture:browse. The high values highlight significant challenges in both wet and dry season in terms of ME availability(Gabler and Heinrichs, 2003; Yeom et al., 2002) and the need to identify supplements with higher ME values.

The seemingly favourable nutritional profile of *Viscum* spp. (ME 11.5 MJ/kg) suggests it could be an effective supplement to improve nutrition, particularly during the dry season. This is further supported by anthelmintic properties reported elsewhere (Madibela et al., 2010; Moncho et al., 2012). Madibela and Jansen (2003) reported no adverse effects of *Viscum* spp. supplementation, however research is limited and, especially at higher concentrations, caution should be taken, and long-term research conducted. Forage preservation may be necessary to facilitate supplementation, however this is not a common practice in the region, leaving animal nutrition at the mercy of the environment, particularly weather. Creating stocks of persevered forages could allow farmers to withstand periods of poor forage availability/nutrition and other adverse events (e.g., drought and disease). However, forage preservation is complex, and farmers will have varying capacity to do this. Perhaps community driven and cooperative schemes could be better placed to achieve this, with technical support.

The nutritional composition of supplements and other feeds collected within the study were reported and adds to existing literature and resources such as Feedipedia. However, further work is needed as the external validity of our data, and indeed many of the Feedipedia current resources, is limited in that we were unable to quantify variation in nutrition of those feeds across time and space and their availability may vary greatly between locations. However, this does highlight potential intervention opportunities that may warrant further investigation, especially as many of these identified feeds are underutilised or waste by-products. For example, sunflower heads had an ME content of 9.3 MJ kg^-1^ which is relatively high compared to the dry season feed-baskets formulated, highlighting how they may be able to act as an effective supplement. Strikingly, *Acacia giraffe* seeds had high levels of CP (26.9%) and ME (13.5 MJ kg^-1^) which could not only supplement shortfalls in nutrition but bolster nutrition even at the best of times to increase performance. However, toxicological screening is recommended to ensure safety for broad consumption. In addition, caution must be adopted as estimating ME by equations has limitations and the accuracy of estimates may reduce when applied to uncommon feeds which were not used in the development of the original equation.

This study focussed on macronutrients (fibre, protein, and energy); however, micronutrient (minerals and vitamins) balances are also important. Notably, phosphorus availability is an issue in Botswana and much of SSA (Setshogo et al., 2011; Verde and Matusso, 2014). Further investigation of these diets would help to ensure micronutrient requirements are best met and enable targeted intervention of deficiencies. For the time being, allowing goats some freedom to forage and ensuring they have a diet comprising a variety of forages, may be the best way to mitigate potential micronutrient deficiency risk. Future studies thus need to investigate the interaction effects and practicality of different feeds under farmer led systems.

## 5 Conclusion

Natural pastures and browse play, and will continue to play, an important role in the nutrition and feeding systems of goats in Africa. These feed resources are subjected to seasonal fluctuations, that limit their capacity to cover livestock requirements. Indeed, feed budgets from basal diet resources (pasture and browse) in SSA show a deficit, especially in terms of ME. Therefore, supplementation must be utilised to ensure acceptable production levels and health. Here we discussed several potential feeds and suggestions were made as to how they could be used to develop feed baskets in the dry and wet seasons for goats. Forages in Botswana were found to be nutritionally diverse, not just between species, but also across time (season) and space (soil type). Whilst optimising nutrition is important all year around, the greatest gains appear possible during the dry season, when supplementation can both improve the nutritional quality of feed-baskets, in addition to making up for potential shortfalls in overall forage availability. However, all supplementation is not equal and there are distinct differences in nutrition, availability, and intake rates. Supplementation with *Viscum* spp. appears to hold significant potential and requires further and detailed study.

## 6 Acknowledgements

We are sincerely grateful to the small holder farmers who were involved in this study for their time, patience, knowledge, and willingness to participate. We acknowledge the Ministry of Agriculture, Department of Veterinary Services, Botswana for support. We also acknowledge the outstanding individual contribution of the small livestock Technical Assistant for Serowe, Mr. Ntebolang Ditsela.

Thanks are also extended to colleagues at BIUST (Botswana), LUANAR (Malawi), the University of Pretoria (South-Africa), Rothamsted Research (UK), Harper Adams University (UK), and Queen’s University Belfast (UK), for advice assistance and support during the course of this study.

## 8 Ethical statement

This work was given ethical approval by the Animal Research Ethics Committee of the Directorate of Research and Development, Botswana International University of Science and Technology.

